# The BTK-DDX41 axis of the STING pathway is activated during cytomegalovirus lytic infection

**DOI:** 10.1101/2024.10.03.616496

**Authors:** Vargab Baruah, Christine M. O’Connor

**Author notes:** (V.B.).

## Abstract

The innate immune response is the first line of defense against invading pathogens, including the betaherpesvirus, human cytomegalovirus (CMV). The host’s innate response acts as the first line of defense, and CMV, like other viruses, has consequently evolved multiple mechanisms to manipulate host interferon (IFN) responses. DEAD-box Helicase 41 (DDX41) is an intracellular dsDNA sensor that, upon activation by Bruton’s tyrosine kinase (BTK), triggers type I IFN production through the Stimulator of Interferon Genes (STING) signaling pathway. Here, we show the activation of this signaling pathway during lytic CMV infection, wherein BTK, DDX41, and STING are activated through tyrosine phosphorylation, and both DDX41 and BTK interact with STING. Further, CMV infection re-localizes DDX41 from the nucleus to the cytoplasm, where it localizes to the perinuclear virus assembly compartment (vAC). Here, DDX41 phosphorylation is attenuated, suggesting cytoplasmic redistribution leads to a less active or inactive form. Additionally, DDX41 co-localizes in the vAC with the CMV tegument proteins, pp65 and pp71, each of which interact with DDX41 in immunoprecipitation assays. We further demonstrate the protective role of this signaling pathway, as treatment with the BTK inhibitor, orelabrutinib, attenuates DDX41 phosphorylation/activation and supports increased expression of viral proteins and virus replication. In sum, our work highlights the important role of BTK-DDX41-STING signaling in the innate immune response against CMV, which the virus subverts by attenuating its cytoplasmic activity, thereby diverting it from its typically protective function.

**Importance:** Human cytomegalovirus (CMV) is a ubiquitous pathogen that poses a significant threat to immunocompromised individuals, highlighting the critical role of innate immunity in controlling this viral infection. Despite extensive research, the complex mechanisms underlying innate immunity against CMV and the virus’s strategies for evading immune detection remain only partially understood. This study identifies the activation of the cellular BTK-DDX41-STING innate signaling axis during lytic CMV infection, which ultimately results in protective interferon responses. Our findings show that CMV infection triggers the cytoplasmic redistribution of the cellular protein, DDX41, leading to reduced phosphorylation and activity, thereby undermining its protective function. Additionally, pharmacological inhibition of BTK enhances viral protein expression and replication, highlighting the importance of this pathway in immune defense. Our work identifies BTK- and DDX41-dependent STING signaling as important for innate immune responses against CMV and further advances our understanding of CMV’s manipulation of these responses.

## Introduction

Human cytomegalovirus (CMV), a widespread beta-herpesvirus, infects 40% to 60% of adults in developed countries, with seropositivity nearing 100% in developing nations (1). Following primary CMV infection, the virus enters a latent phase within myeloid cells, characterized by viral genome maintenance and suppressed infectious virus production. Latent CMV can intermittently reactivate under certain conditions of immune system dysfunction and cellular differentiation, resulting in the lytic replication of infectious virus (2). While primary infection and reactivation events are typically asymptomatic, they can be severe in immunocompromised, immunosuppressed, or immuno-naïve individuals, potentially causing serious organ-specific diseases or even death. A substantial proportion of the population, even in developed countries, is immunologically vulnerable. In the US, about 3% of adults are immunocompromised, and 6.6% are immunosuppressed due to health conditions or their treatments (3, 4). Among immuno-naïve neonates, approximately 1 in 200 babies has congenital CMV infection (5), and up to 15% of these infants develop long-term sequelae (6). These data underscore the need to more completely understand the viral and cellular factors that support CMV replication.

The severity and outcome of CMV infection largely depend on the early immune events, when the host’s innate immunity plays a crucial role (7). It can prevent the establishment of infection by overwhelming CMV upon initial contact or confine the virus to the latent phase, preventing progression to the lytic phase. Early immune responses often involve recognition of pathogen-associated molecular patterns (PAMPs) – conserved molecular signatures found on pathogens - by the host’s pattern recognition receptors (PRRs), which heavily influence the subsequent immune strategy. CMV’s double-stranded DNA (dsDNA) genome is detected by dsDNA-sensing PRRs, triggering an immune signaling cascade that results in the production of anti-viral interferons (IFNs) and interferon-stimulated genes (ISGs) (8). Understanding these key immune responses at the molecular level is crucial, particularly in the absence of a vaccine, to determine how CMV might subvert these mechanisms as it infects the host.

Activation of key components of the innate immune response is intricately linked to the CMV life cycle, with different viral components targeted based on their abundance and accessibility. CMV dsDNA can bind cytosolic DNA receptors such as cyclic GMP-AMP synthase (cGAS) and interferon-γ inducible protein 16 (IFI16). This DNA sensing is followed by activation of stimulator of interferon genes (STING; also known as MITA and ERIS), a key mediator in subsequent downstream immune signaling. STING activation leads to the recruitment and activation of TANK-binding kinase 1 (TBK1), which in turn phosphorylates interferon regulatory factor 3 (IRF3). This activated IRF3 then translocates to the nucleus, where it drives the expression of type I interferons (IFN) and other inflammatory cytokines, thereby mounting a defense against the invading virus (9).

Host-driven immune responses against CMV are complex since CMV has evolved impressively sophisticated methods of manipulating host factors. For instance, CMV’s tegument protein pp65 and late protein pUL31 bind cGAS, inhibiting its activity (10, 11). Additionally, the membrane protein pUL42 also binds cGAS, blocking both its DNA binding and its ability to oligomerize/activate (12). Similar observations are reported for IFI16, where pp65 binds IFI16 and inhibits its DNA sensing by preventing its aggregation and signal amplification (13, 14). CMV also modulates restriction factors downstream of DNA sensing. The viral glycoprotein, US9, the tegument protein, pp71, and the latency-associated protein, pUL138, each bind STING and impair immune signaling (15–17). In addition to directly inhibiting immune signaling through binding, CMV also subverts the immune response by misdirecting the localization of restriction factors. For instance, while nuclear IFI16 binds CMV DNA early in infection, during later stages, it is phosphorylated by pUL97 and mis-localized to the cytoplasmic vAC (18). While these studies highlight the intricate and diverse mechanisms the host employs against CMV, as well as those the virus uses to counter them, our understanding of the array of innate immune sensors triggered by CMV is likely incomplete.

DEAD-box helicase 41 (DDX41) is a helicase with a conserved Asp-Glu-Ala-Asp (DEAD) motif that functions as a DNA sensor (19). Similar to other DNA sensors, such as cGAS and IFI16, DDX41 activity is regulated by tyrosine phosphorylation. Lee at al. demonstrated that Bruton’s tyrosine kinase (BTK) phosphorylates DDX41 enhancing its binding to adenovirus dsDNA and STING (20). Additionally, herpes simplex virus type 1 (HSV-1) infection results in upregulation of DDX41, upon which it interacts with cGAS, activates the STING signaling pathway, and promotes inflammatory cytokine production (19). However, the role of the BTK-DDX41-STING signaling axis in CMV infection remains unknown. Given that DDX41 shares many downstream effectors with cGAS and IFI16, and that CMV has a dsDNA genome, we posited that this signaling pathway could function as part of the innate immune response against CMV.

Here, we show that the BTK-DDX41-STING signaling axis is activated during lytic CMV infection, involving tyrosine phosphorylation of the restriction factors, and interaction of STING with DDX41 and BTK. Following infection, DDX41 re-localizes from the nucleus to the cytoplasm, where it co-localizes with the tegument protein, pp65, in the perinuclear vAC. Additionally, cytoplasmic-localized DDX41 displays reduced phosphorylation, indicating that its cytoplasmic redistribution leads to a less active form of the protein. Finally, we demonstrate that inhibiting BTK with orelabrutinib reduces phosphorylation and activation of BTK and DDX41, which increases viral protein expression and replication. Our work reveals an important role of the BTK-DDX41-STING pathway in the innate immune response to CMV and demonstrates a mechanism by which the virus disrupts this response.

## Materials and Methods

### Cells and Viruses

Primary newborn human foreskin fibroblasts (NuFF-1, passage 13-28; GlobalStem, Rockville, MD, USA) were cultured in Dulbecco’s modified Eagle medium (DMEM) with 10% fetal bovine serum (FBS), 2 mM L-glutamine, 0.1 mM non-essential amino acids, 10 mM HEPES, and 100 U/ml each of penicillin and streptomycin. The cells were maintained at 37°C with 5% CO_2_.

The bacterial artificial chromosome (BAC)-derived viruses, TB40/E*mCherry* (WT) (21) and TB40/E*mCherry*-UL99eGFP (UL99*gfp*) (22, 23) were previously characterized. Virus stocks were prepared as previously described (24). In brief, p0 stocks were generated by transfecting BAC DNA into naïve NuFF-1 cells. When 100% cytopathic effect (CPE) was observed, p1 stocks were expanded on naïve NuFF-1 cells and harvested when 100% CPE was observed. Virus was purified through a 20% sorbitol (VWR) cushion, resuspended 1:1 in XVIVO-15 (Lonza) and 3% bovine serum albumin (BSA), aliquoted, flash frozen in liquid nitrogen, and stored at −80°C. Virus stocks were titered on naïve NuFF-1 cells by 50% tissue culture infectious dose (TCID_50_) assay.

Cells were infected at the multiplicity of infection (MOI) and for the duration indicated in the text. Cells were infected in DMEM containing viral inoculum for 1 h at 37°C, with gentle agitation every 15 minutes (min). The cells were washed twice with 1X phosphate-buffered saline (PBS) and replenished with fresh DMEM at 37°C.

### Protein, RNA, and DNA Analyses

For protein analyses, cells were treated and harvested at the times indicated in the text and/or Figure Legends. Where indicated, cells were transfected with 10 µg/ml of either poly(dA:dT) interferon stimulatory DNA (ISD), or control ISD using Lipofectamine 2000. In all cases, cells were lysed on ice for 1 hour (h) using radioimmunoprecipitation assay (RIPA) buffer (1.0% NP-40; 1.0% sodium deoxycholate; 0.1% SDS; 0.15 M NaCl; 0.01 M NaPO4, pH 7.2; 2 mM EDTA), with vortexing every 15 min. For protein isolation in nuclear/cytoplasmic fractions, cell pellets were resuspended in lysis buffer (10 mM HEPES, pH 7.5; 5 mM MgCl2; 15 mM KCl; 1 mM PMSF) and subjected to three freeze-thaw cycles in liquid nitrogen. The resulting suspension was passed through a 27G needle 15 times. The lysed cells were layered over 50% sucrose (w/v) and centrifuged; the supernatant collected as the cytoplasmic fraction. The pellet containing nuclei was resuspended in lysis buffer and sonicated three times. After centrifugation, the supernatant was collected as the nuclear fraction. In all cases, protein concentrations were measured by Bradford assay with Protein Assay Dye Reagent Concentrate (Bio-Rad). Samples were denatured at 95°C for 10 min. Equal protein concentrations of the samples were separated by SDS-PAGE and transferred to 0.45 µm nitrocellulose membranes (Amersham) using a semi-dry transfer method. The following antibodies were used at the specified concentrations: anti-BTK (Cell Signaling Technology [CST], 1:5,000), anti-STING (CST, 1:5,000; MilliporeSigma, 1:5,000), anti-p-STING (CST, 1:1000), anti-DDX41 (CST, 1:5,000), anti-TBK (CST, 1:5,000), anti-p-TBK (CST, 1:1000), anti-CMV IE1 (clone 1B12, 1:100, (25)), anti-CMV UL44 (Virusys, 1:2500), anti-CMV pp65 (clone 8A8, 1:50 (26)), anti-CMV pp71, anti-tubulin (MilliporeSigma, 1:5000), anti-H3 (Abcam, 1:5000), anti-beta-actin-peroxidase (MilliporeSigma, 1:10,000), goat-anti-mouse horseradish peroxidase (HRP) secondary and goat-anti-rabbit horseradish peroxidase (HRP) secondary (both from Jackson ImmunoResearch Labs, 1:10,000). Relative protein levels were quantified by densitometry using ImageJ (27).

To quantify transcripts, total RNA was isolated and purified using the High Pure RNA Isolation kit (Roche) in accordance to the manufacturer’s protocol. Equal RNA concentrations (1.0 μg) were then used to generate cDNA with random hexamer primers and SYBR™ Green I Reagents (ThermoFisher Scientific), and equal volumes of cDNA were used for quantitative polymerase chain reaction (qPCR). The following primers were used for transcript quantification: *IFNB*, forward – 5’-CTTGGATTCCTACAAAGAAGCAGC-3’ and reverse – 5’-TCCTCCTTCTGGAACTGCTGCA - 3’; *UL123*, forward – 5’-GCCTTCCCTAAGACCACCAAT-3’ and reverse – 5’-ATTTTCTGGGCATAAGCCATAATC-3’; *GAPDH*, forward – 5’-CTGTTGCTGTAGCCAAATTCGT-3’ and reverse – 5’-ACCCACTCCTCCACCTTTGAC-3’. Samples were analyzed in triplicate using a 96-well-format CFX Connect Real Time PCR machine (Bio-Rad), and expresion quantified relative to host *GAPDH* expression.

To quantify CMV genomic DNA, NuFF-1 fibroblasts were infected (MOI = 0.5 TCID_50_/cell) with WT virus. At each indicated time point, cells were harvested and samples were treated with RNaseA (100 µg/mL) for 1 h at 37°C to remove RNA. DNA was isolated by phenol-chloroform-isoamyl alcohol extraction, followed by ethanol precipitation and resuspension in 10 mM Tris-HCl, pH 8.0 with 0.1 mM EDTA. Equal sample volume was quantified by qPCR using the following primers: viral UL69, forward – 5’-GGCTGATGATCTTGCGGGAA-3’ and reverse – 5’-CGAGAGTCTACGTCTGGCAC-3’; cellular MDM2, forward – 5’-CCCCTTCCATCACATTGCA-3’ and reverse – 5’-AGTTTGGCTTTCTCAGAGATTTCC-3’. UL69 and MDM2 abundance was quantified against a viral BAC standard (TB40/E-BACstandard), which contains an MDM2 gene fragment (28). All samples were analyzed in triplicate, then normalized to cellular MDM2.

### Immunofluorescence Assays

Cells were cultured on coverslips in 6-well tissue culture plates (3 × 10^5^ cells per well) and infected (MOI = 0.5 TCID_50_/cell) as described above. After 96 hpi, cells were washed three times with 1X PBS, fixed with 2% paraformaldehyde for 15 min at 37°C, and then permeabilized with 0.1% Triton X-100 for 15 min at room temperature. The cells were washed three times with 0.25% (w/v) saponin and subsequently blocked with 1X PBS containing 10% (v/v) human serum (Sigma Aldrich) and 0.25% (w/v) saponin. Cells were incubated with primary antibodies against STING, BTK, phospho-BTK-Y551 (all from Cell Signaling Technology, 1:1000), DDX41 (Cell Signaling Technology, 1:1,000), and STING (MilliporeSigma, 1:1,000) in 10% (v/v) human serum and 0.25% (w/v) saponin for 1 h at room temperature. Following this, the cells were washed three times with 0.25% (w/v) saponin and incubated with Alexa-488-conjugated anti-mouse and Alexa-647-conjugated anti-rabbit secondary antibodies (both from Thermo Fisher, 1:1,000) for 1 h at room temperature, shielded from light. Finally, coverslips were mounted with FluorSave reagent (MilliporeSigma) onto slides and visualized using a Leica SP8 inverted confocal microscope with a 63x oil immersion objective. Colocalization in immunofluorescent images was quantified using Volocity software. To assess colocalization of DDX41 or TBK1 with STING, we calculated the percentage of mCherry-positive, CMV-infected cells that exhibited overlap between the Volocity-defined compartments of each protein and STING. Quantification was performed on three fields of view per condition, and the resulting percentages were plotted as bar graphs.

### Co-immunoprecipitation

Cells were lysed in 1.0 ml of immunoprecipitation (IP) buffer [25 mM Tris, pH 7.4; 1 mM MgCl₂; 2 mM EDTA; 1.0% Triton-X 100; 150 mM NaCl; Complete Protease Inhibitor Cocktail (Roche)] on ice for 2 h. Protein G beads (100 μl; Protein G-sepharose Fast Flow, MilliporeSigma) were washed twice with IP buffer (500 x *g*, 5 min), blocked for 1 h in 5.0% BSA in IP buffer, and then resuspended in 1.0 ml IP buffer. The samples were clarified by centrifugation (12,000 x *g*, 10 min, 4°C), and 50 μl of the sample (10%) was retained as the “input” control. The remaining sample was divided into two aliquots, each incubated overnight at 4°C with protein G beads and either 2 μg of the specific precipitating antibody or an isotype-matched IgG control antibody. The next day, the samples were centrifuged at 500 x *g* for 5 min, and 50 μl was reserved as the “unbound” sample. The samples were then washed four times with IP wash buffer [25 mM Tris, pH 7.4; 10 mM MgCl₂; 2 mM EDTA; 0.1% Triton-X 100], with 50 μL retained after washes 1 and 4. After the final wash, proteins in all samples were denatured using 2X SDS and heating at 95°C for 10 min. The samples were then separated by SDS-PAGE and transferred to nitrocellulose as described above.

### Toxicity Assay

NuFF-1 cells (1.75 x 10^4^ cells/well) were plated onto 96-well opaque white CulturPlate (Perkin Elmer) and allowed to adhere overnight. Cells were then treated with orelabrutinib (GLPBio) in nine two-fold dilutions (0.078 to 20 µM) for 24 h, followed by three washes with 1X PBS. Cell viability was assessed after 2 and 5 days (i.e., 3 and 6 days post-treatment) using CellTiter-Glo (Promega) according to the manufacturer’s guidelines, and the plates were read with a Cytation 5 (Agilent) plate reader. DMSO (equivalent v/v) served as the diluent and vehicle control, while puromycin (ranging from 20 µM to 0.08 µM) was used as a positive control.

### Quantification of viral lytic replication

NuFF-1 cells were treated with 0.5 µM orelabrutinib (GLPBio) prior to infection with TB40/E*mCherry* (MOI = 0.01 TCID_50_/cell) for 1 h, with gentle agitation every 15 min. After adsorption, cells were washed three times with 1X PBS, and fresh media (V_f_ = 2.0 ml) containing 0.5 µM Orelabrutinib was added. Cell-free virus (350 µl) was collected over a 16 day (d) timecourse, and 250 µl of fresh media with 100 µl of 10 µM orelabrutinib was added to restore the total volume to 2 ml. DMSO (equivalent v/v) served as the diluent and vehicle control. Cell-associated virus was harvested at the final time point (16 days post-infection; dpi) by lysing the cells through three freeze-thaw cycles. The clarified supernatant was then used to determine the cell-assocated viral titer. Cell-free and cell-associated virus were quantified by TCID_50_ assay on naïve NuFF-1 fibroblasts.

## Results

### BTK-DDX41-STING signaling is activated during lytic CMV infection

Emerging data reveal an antiviral role for DDX41 (19, 29–31), though whether DDX41 functions as a restriction factor during CMV infection remains unclear. To begin to address this, we first assessed the expression and activation of restriction factors associated with the BTK-DDX41-STING signaling pathway during lytic CMV replication. We mock- or lytically-infected NuFF-1 fibroblasts with TB40/E*mCherry* (wild type; WT) over a 96 hour (h) time course, after which we measured protein BTK, STING, DDX41 and TBK1 abundance and phosphorylation, which is essential for activation of these restriction factors (20, 32–34). Additionally, we probed for the viral protein, IE1. As a control for BTK and DDX41 activation, we transfected parallel cultures with poly(dA:dT) (29). Since there is no phospho-specific antibody for DDX41, and the phospho-BTK antibody is ineffective in immunoblot assays, we performed immunoprecipitation with a pan-phospho-tyrosine antibody, followed by immunoblot using antibodies directed at total forms of either BTK or DDX41. Our data indicate BTK is activated by 6 h post-infection (hpi), which is increased through 96 hpi (**Fig. 1A**). Similarly, DDX41 is also activated at 6 hpi, with sustained increased phosphorylation through the 96 h time point. Additionally, total DDX41 is also increased over the time course, relative to mock-infected cultures (**Fig. 1B**). Levels of both total and phosphorylated forms of BTK and DDX41 were comparable to those induced by poly(dA:dT) transfection over a similar time course, although poly(dA:dT) transfection triggered an earlier and more rapidly peaked response (**Fig. 1A, 1B**). We also observed an increase in phospho-forms of STING and TBK1 to levels consistent with their activation following interferon stimulatory DNA (ISD) transfection (**Fig. 1C**), which is a positive control for STING-TBK1 activation (35). Finally, we quantified mRNA expression of *IFNB*, which encodes IFN-β, a protein upregulated in response to innate immune signals. Infection induces a time-dependent increase in *IFNB* transcription, which peaks at 3 hpi and is sustained throughout the time course (**Fig. 1D**). We consistently observed increased activation of BTK and DDX41 at 96 hpi, with sustained activation of STING and TBK1 through this time point. Overall, these data indicate CMV induces both the abundance and phosphorylated activation of BTK, DDX41, STING, and TBK1, which results in increased *IFNB* expression in infected fibroblasts.

**Figure 1.**
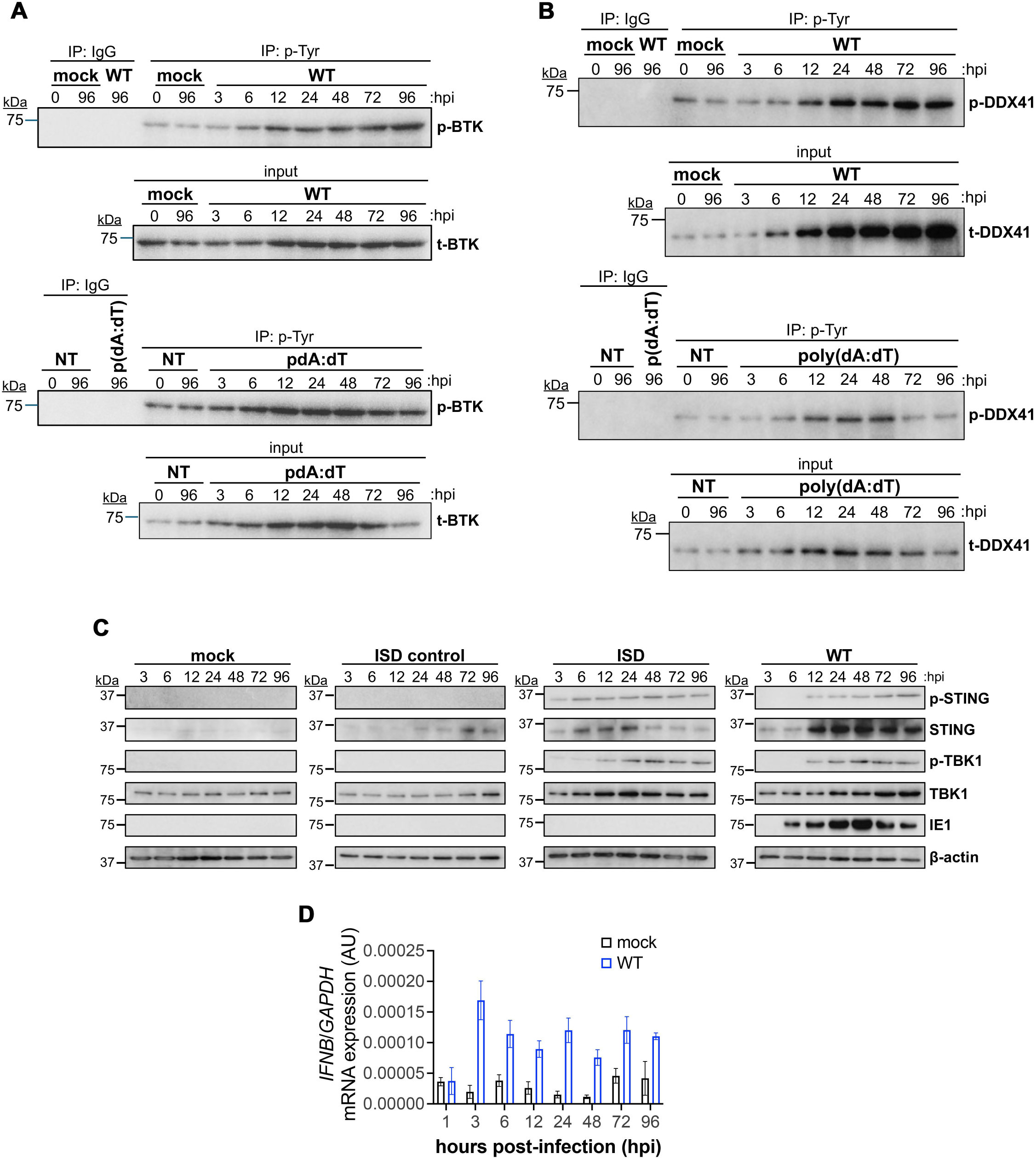
CMV infection induces a time-dependent increase in BTK, DDX41, STING, TBK1 abundance and activation, as well as *IFNB* mRNA expression. (**A, B**) NuFF-1 fibroblasts were mock- or WT-infected (MOI = 0.5 TCID_50_/cell) or transfected with 10 µg/mL poly(dA:dT). At the indicated times post-infection or -transfection, cells were immunoprecipitated (IP) with antibodies specific for phosphorylated-tyrosine (p-Tyr) or IgG. Immunoprecipitates were then probed for BTK and DDX41 by western blot (WB). Input for the IPs are also shown. N = 3; representative blots shown. (**C**) NuFF-1 fibroblasts were mock- or WT-infected (MOI = 0.5 TCID_50_/cell) or transfected with 10 µg/mL interferon stimulatory DNA (ISD) or ISD control DNA. At the indicated times post-infection or -transfection, cells were collected and equal concentration (30 µg) of lysates were probed for phosphorylated (p) and total forms of the cellular proteins, STING and TBK1, as well as the viral protein, IE1. Cellular β-actin was probed as a loading control. N = 3, representative blots shown. (**E**) Total RNA was isolated at the indicated times post-infection, and *IFNB* was measured by RT-qPCR and plotted in arbitrary units (AU) as ΔΔCt relative to cellular *GAPDH*. N = 3 biological replicates, each with technical triplicates; representative experiment shown.

To investigate the signaling mechanism and confirm pathway activation, we used immunofluorescence assays (IFA) to examine colocalization and co-immunoprecipitation (co-IP) to assess interactions among restriction factors, with a particular focus on STING. Previous studies have shown STING physically interacts with many of its signaling partners, such as TBK1 and IRF3, bringing them into close proximity for phosphorylation and subsequent activation (36). We mock- or WT-infected NuFF-1 fibroblasts for 96 h and visualized colocalization of each restriction factor (BTK, pBTK-Y551, DDX41 and TBK) with STING by immunoflourescence. We found BTK (**Fig. 2A**), its activated form pBTK-Y551 (**Fig. 2B**), DDX41 (**Fig. 2C**), and TBK1 (**Fig. 2D**) each colocalize with STING at the perinuclear region of the infected cell. We quantified the colocalization of STING with upstream DDX41 and downstream TBK1 in infected cells expressing mCherry between 24 and 96 hpi and found consistent colocalization during this period in approximately 40–50% of cells (**Fig. 2C, 2D**). Quantification in mCherry-positive cells was not possible between 3-12 hpi due to the absence of mCherry expression (**Fig. S1**). Further, we confirmed the interaction between STING and BTK (**Figs. 3A, S2A**), as well as STING and DDX41 (**Figs. 3B, S2B**) by co-IP, consistent with previous findings (20, 37). We also confirmed CMV infection by IE1 protein abundance (**Fig. S2C,D**). Collectively, our data reveal BTK-DDX41-STING signaling is activated following WT infection,.

**Figure 2.**
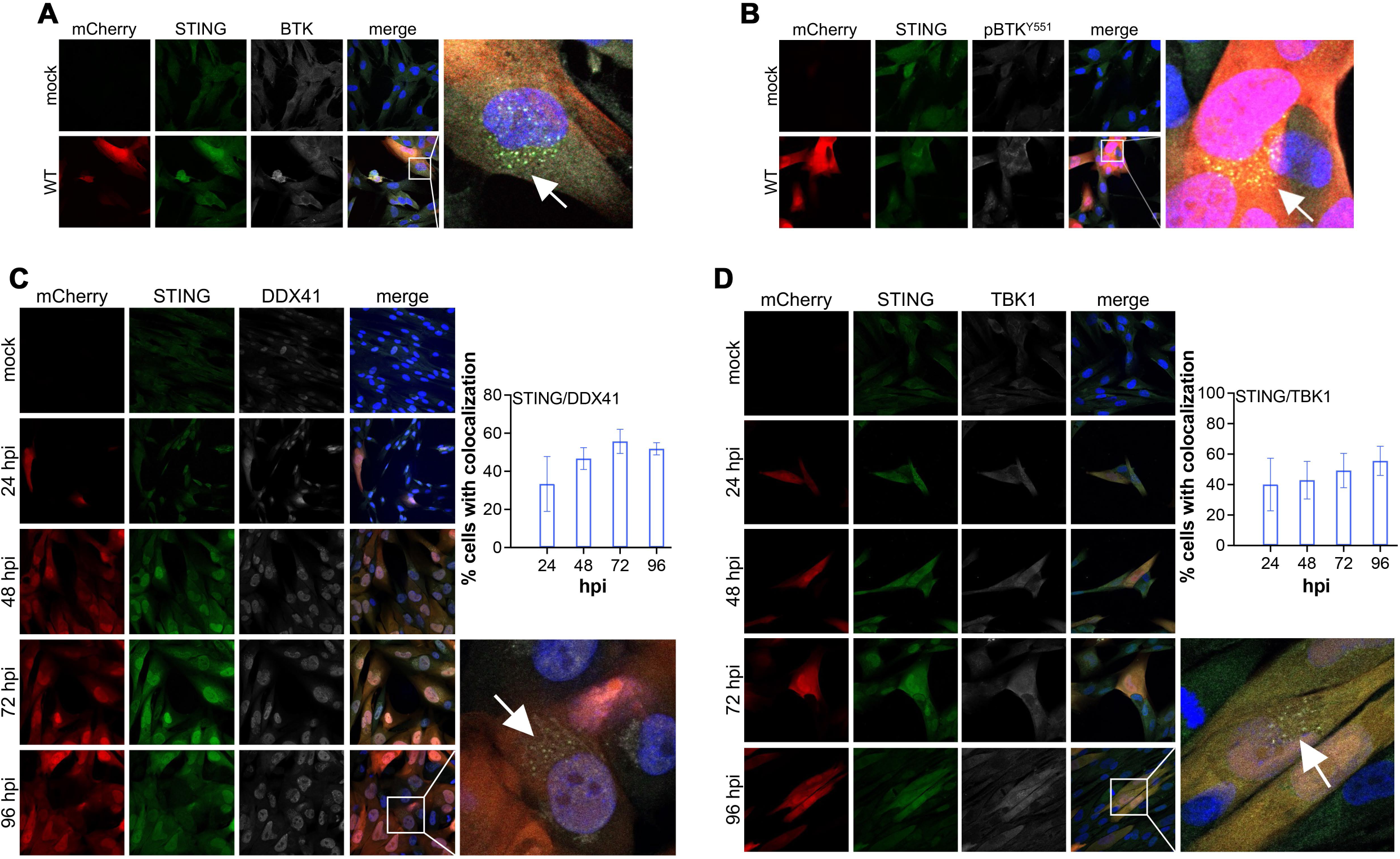
BTK, DDX41, and TBK1 co-localize with STING during CMV infection. NuFF-1 fibroblasts were mock- or WT-infected (MOI = 0.5 TCID_50_/cell), after which cells were processed for immunofluorescence assay (IFA) at (**A, B**) 96 hpi or (**C,D**) over a time course of infection. The early time points (3-12 hpi) for the data in (**C,D**) are shown in Fig. S1. Cells were then probed with antibodies directed at: (**A**) BTK, (**B**) pBTK-Y551, (**C**) DDX41, or (**D**) TBK1 (each shown in white), as well as (**A-D**) STING (green). (**C,D**; inset graph) The percentage of infected, mCherry-positive cells displaying (**C**) STING and DDX41 or (**D**) STING and TBK1 colocalization was quantified using Volocity. mCherry (red) is a marker of lytic infection; nuclei were visualized with DAPI (blue). Inset in the WT merge panel is enlarged at right. Images were acquired using 63x objective. N = 3; representative images shown.

**Figure 3.**
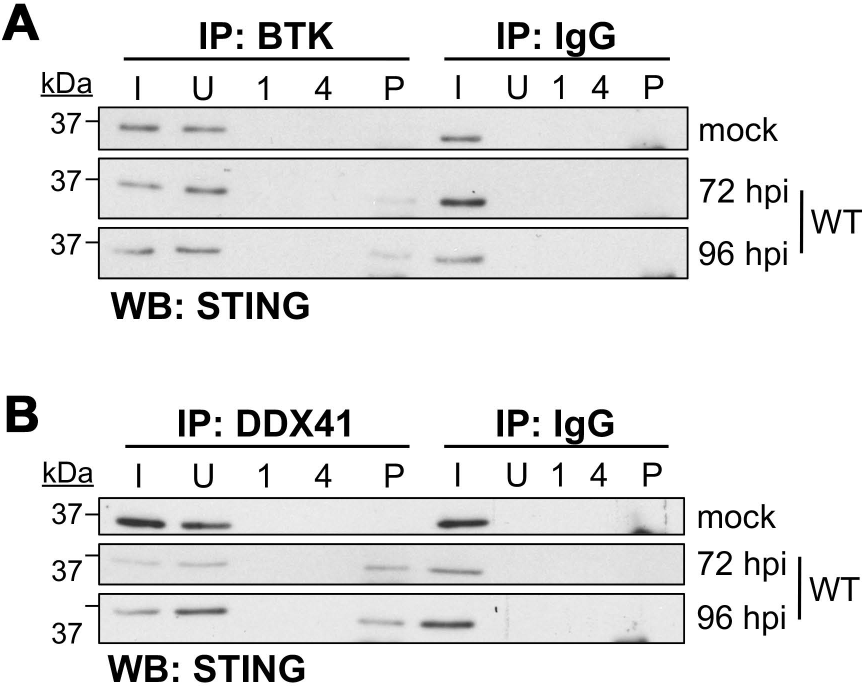
BTK and DDX41 interact with STING during CMV infection. NuFF-1 fibroblasts were mock- or WT-infected (MOI = 0.5 TCID_50_/cell) for the indicated times. Cells were then harvested, and immunoprecipitated (IP) with antibodies specific for (**A**) BTK, (**B**) DDX41, or (**A,B**) IgG. Immunoprecipitates were then probed for STING by western blot (WB). I, input; U, unbound; 1, wash #1; 4, wash #4; P, IP sample. N = 3; representative blots shown.

### CMV redistributes a fraction of DDX41 to the cytoplasm

Our data reveal CMV infection results in distinct changes in the subcellular localization of the restriction factors, including DDX41 (**Fig. 2**). In the absence of infection (mock control), DDX41 primarily displays nuclear localization; however, CMV infection results in the progressive migration of this host restriction factor to the perinuclear region as early as 24 hpi. DDX41 cytoplasmic localization increases as infection progresses and is observed in about 50-60% of all infected cells at 72 and 96 hpi (**Fig. 2C**). This is consistent with previous work revealing DNA stimulation triggers DDX41 relocalization (19). Further, the CMV-induced perinuclear relocalization of DDX41 suggested to us that this host protein may become co-localized within the CMV vAC. To test this, we infected fibroblasts with a recombinant virus, TB40/E*mCherry*-UL99*gfp*, which contains eGFP fused to the carboxy terminus of the *UL99* ORF (22, 23). *UL99* encodes pp28, a viral protein that localizes to the vAC during lytic infection (38), which we also confirmed (**Fig. 4A**). Similar to our data in **Fig. 2C**, we noted a portion of DDX41 relocalizes to the vAC (**Fig. 4A**). Additionally, the other DDX41-associated restriction factors (BTK, STING, TBK1) also relocalize to the vAC following CMV lytic infection (**Fig. S3**).To gain a better understanding of the abundance of DDX41 that was relocalized in response to CMV infection, we analyzed protein lysates derived from nuclear and cytoplasmic fractions of mock- or WT-infected cells over a time course of infection. In mock-infected cells, DDX41 retains its nuclear localization. However, in CMV-infected fibroblasts, a portion of DDX41 is translocated to the cytoplasm as early as 24 hpi, which increases and persists through the time course we evaluated (**Fig. 4B**). We confirmed this cytoplasmic shuttling of DDX41 to the perinuclear vAC between 24 and 96 hpi by IFA for DDX41 and pp65, a viral protein that also localizes to the vAC during lytic infection (**Figs. 4C, S4**). We hypothesized that since DDX41 is a dsDNA sensor, perhaps a portion of DDX41 is retained within the nucleus by the host cell to target the viral genome. To test this, we evaluated whether DDX41 colocalizes with the CMV DNA polymerase subunit, UL44, which we used to identify sites of viral genome replication by IFA. Although a fraction of DDX41 remains in the nucleus during infection, nuclear DDX41 does not specifically colocalize with UL44 (**Fig. 4D**). Taken together, our data indicate DDX41 is relocalized from the nucleus to perinuclear, cytoplasmic regions consistent with the vAC in the majority of infected cells late times in infection.

**Figure 4.**
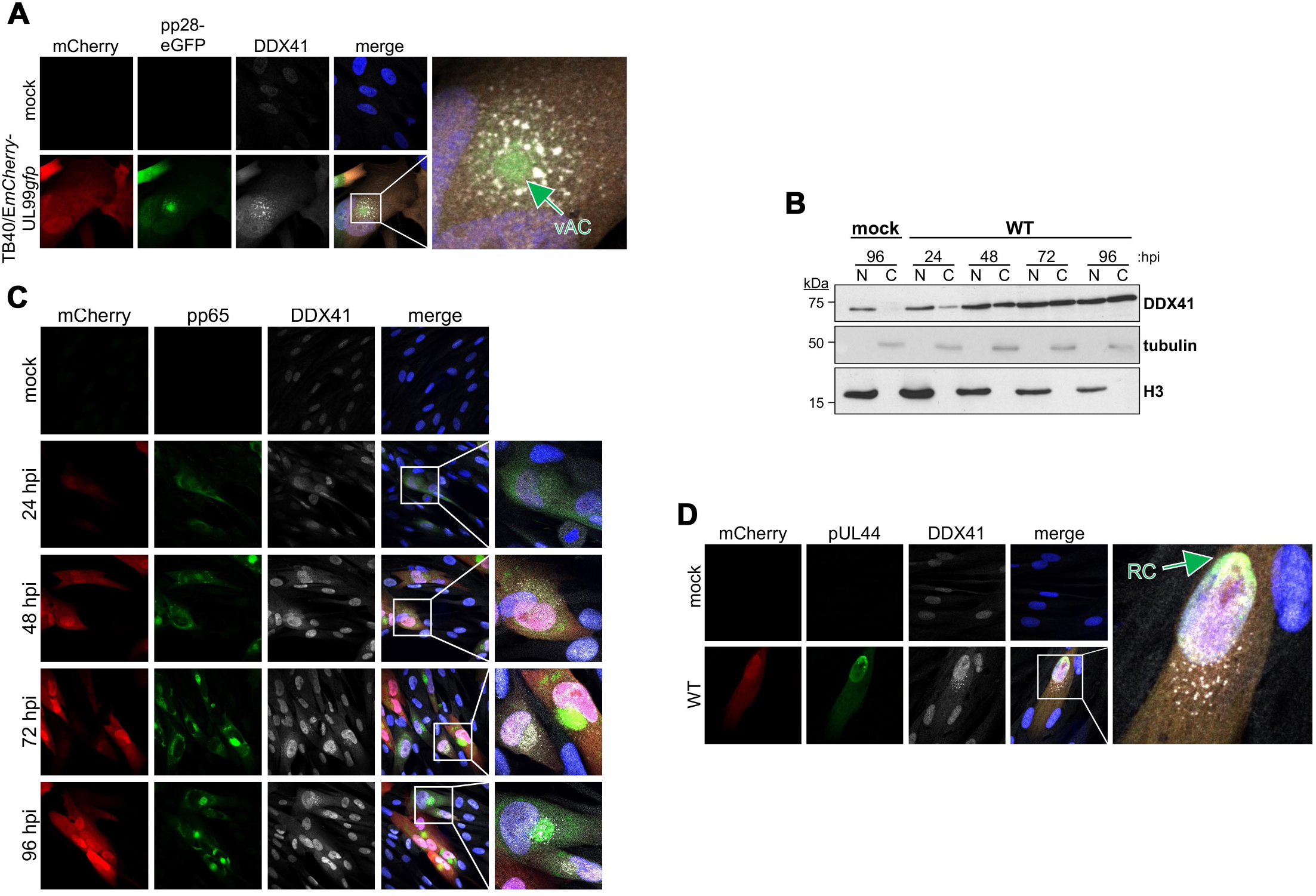
DDX41 is translocated from the nucleus to the perinuclear viral assembly compartment (vAC). (**A**) NuFF-1 fibroblasts were mock- or *UL99gfp*-infected (MOI = 0.5 TCID_50_/cell) for 96 h. Cells were then fixed, permeabilized, and stained for DDX41 (white). mCherry (red) is a marker of lytic infection, pp28-eGFP is shown in green, and nuclei were visualized with DAPI (blue). vAC, viral assembly compartment. (**B**) NuFF-1 fibroblasts were mock- or WT-infected (MOI = 0.5 TCID_50_/cell) for the indicated times. Lysates (20 µg) from nuclear (N) and cytoplasmic (C) fractions were probed for DDX41, tubulin (cytoplasmic marker), and histone H3 (nuclear marker). N = 3; representative blots shown. (**C, D**) NuFF-1 fibroblasts were infected as in (**B**). (**C**) At the indicated times post-infection or (**D**) 96 hpi, cells were fixed, permeabilized, and stained for (**C**) pp65 or (**D**) pUL44 (each shown in green) and DDX41 (white). mCherry (red) is a marker of lytic replication and DAPI (blue) was used to visualize nuclei. Viral replication compartment (RC) is indicated. (**A,C,D**). Inset in the merge panel is enlarged at right. Images were acquired using 63x objective. N = 3; representative images shown.

In addition to relocalizing this antiviral protein, we hypothesized that CMV infection may also result in altered activity of the cytoplasmic portion of DDX41. Thus, we next asked whether perinuclear-localized DDX41 exhibited altered signaling potency compared to its nuclear counterpart. To this end, we analyzed the tyrosine phosphorylation of DDX41 in nuclear and cytoplasmic fractions of cells that were either mock- or WT-infected, first confirming the fractionation using tubulin and H3 as cytoplasmic and nuclear loading controls, respectively (**Fig. 5A**). We then performed immunoprecipitation with a phospho-tyrosine antibody and probed for DDX41 in these fractions. Under mock conditions, DDX41 is localized to the nucleus, consistent with our findings above, where it is phosphorylated. During infection, although total DDX41 is more abundant in the cytoplasm than in the nucleus, the phosphorylated/activated fraction in the cytoplasm is greatly reduced, as indicated by a reduction in the p-Tyr DDX41 pulldown from the cytoplasmic fraction (**Fig. 5B**). Taken together, these data indicate CMV infection induces a relocalization of a fraction of the inactivated form of DDX41 to the cytoplasm.

**Figure 5.**
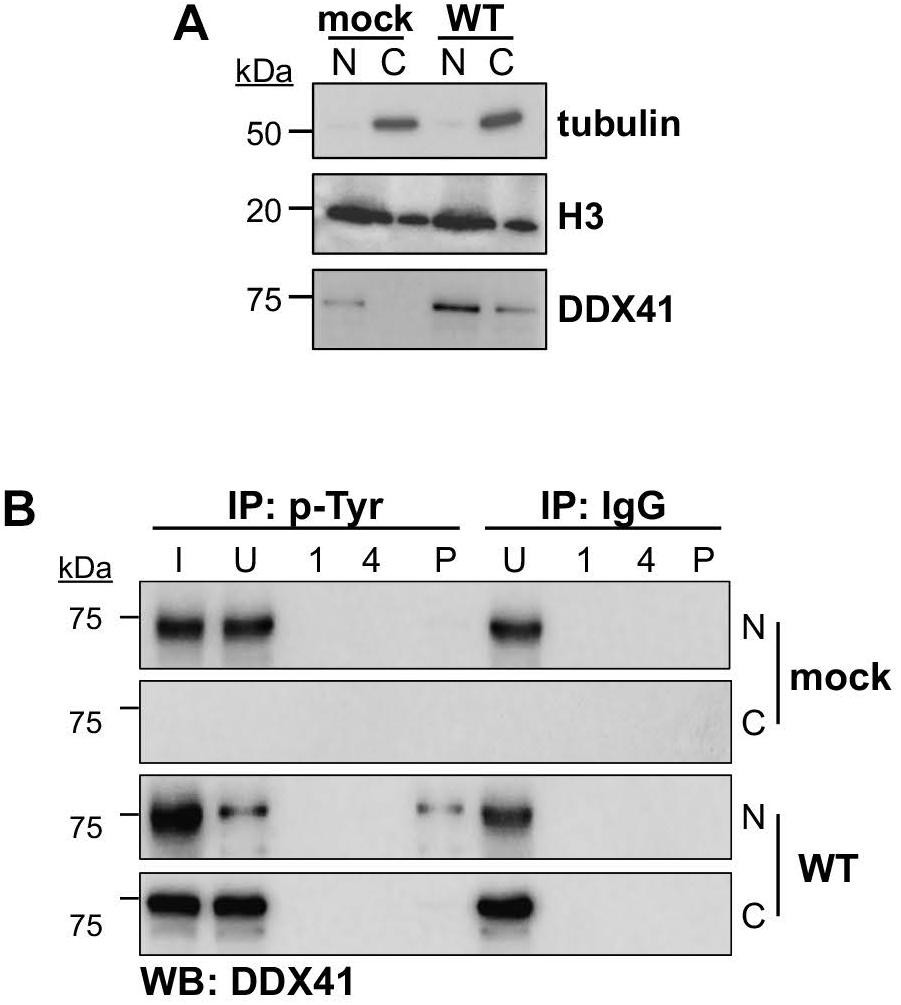
Cytoplasmic DDX41 exhibits attenuated phosphorylation. NuFF-1 fibroblasts were mock- or WT-infected (MOI = 0.5 TCID_50_/cell). At 96 hpi, cells were harvested and nuclear (N) and cytoplasmic (C) fractions isolated. (**A**) Nuclear (N) and cytoplasmic (C) fractions were probed for tubulin (cytoplasmic), histone H3 (nuclear), or DDX41. (**B**) Fractions from (**A**) were then immunoprecipitated (IP) with antibodies specific phosphorylated Tyrosine (p-Tyr) or IgG and were then probed for DDX41 by western blot (WB). I, input; U, unbound; 1, wash #1; 4, wash #4; P, IP sample. N = 3; representative blots shown.

### DDX41 interacts with CMV-encoded tegument proteins

Previous reports have noted a similar CMV-induced perinuclear distribution of the dsDNA sensor, IFI16. At later stages of infection, IFI16 is localized to the vAC, where it interacts with the tegument protein, pp65, to collectively subvert IFI16-mediated DNA sensing (10, 13, 18). Indeed, pp65 also interacts with cGAS, thereby interfering with cGAS-STING interaction and innate sensing activity (10). In parallel, *UL82*-encoded tegument protein, pp71, interacts with host restriction factors, ultimately preventing the proper assembly of the STING-TBK1-IRF complex, whose activity is required for innate sensing via this pathway (39). Based on these observations, as well as our IFAs showing colocalization of DDX41 and pp65 between 24 and 96 hpi (**Fig. 4C**), we hypothesized that a similar interaction between CMV-encoded tegument proteins and DDX41 regulates this host restriction factor’s localization and activity following CMV infection. To this end, we infected fibroblasts and assessed the potential co-localization and interaction between DDX41 and either pp65 or pp71. Our findings reveal DDX41 co-localizes with pp65 (**Fig. 6A**) and pp71 (**Fig. 6B**) in the perinuclear vAC. Further, DDX41 interacts with pp65 (**Fig. 6C**) as well pp71, though detection of this latter interaction is far weaker than the former (**Fig. 6D**). These results suggest that pp65, and to a lesser extent, pp71, interact with DDX41, likely retaining this host restriction factor in the cytoplasmic vAC.

**Figure 6.**
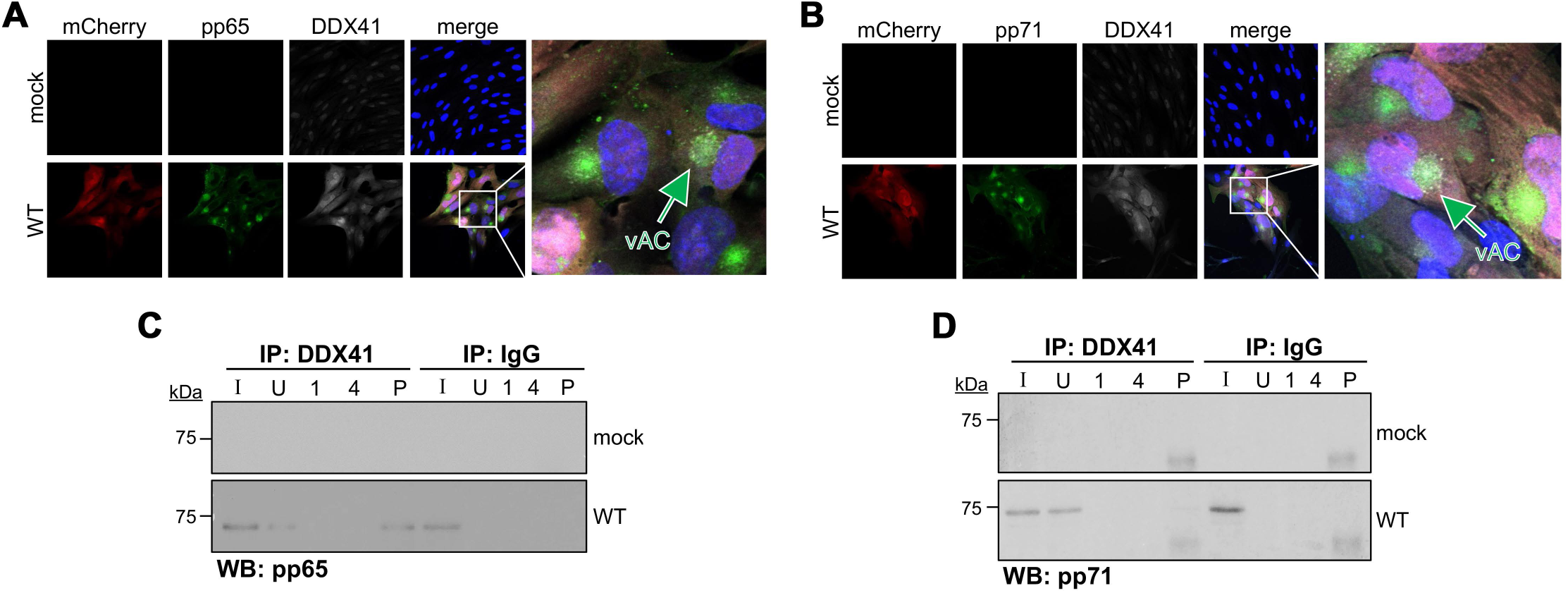
CMV tegument proteins, pp65 and pp71, interact with DDX41 and co-localize with DDX41 in the vAC. NuFF-1 fibroblasts were mock- or WT-infected (MOI = 0.5 TCID_50_/cell) for 96 h. (**A,B**) Cells were fixed, permeabilized and stained for DDX41 (white) and either (**A**) pp65 (green) or (**B**) pp71 (green). (**A,B**) mCherry (red) is shown as a marker of infection, and nuclei were visualized with DAPI (blue). (**C,D**) Cells were then harvested, and proteins immunoprecipitated (IP) with antibodies specific for DDX41 or IgG. Immunoprecipitates were then probed for (**C**) pp65 or (**D**) pp71 by western blot (WB). Samples were also probed for DDX41 by western blot as a control (Fig. S5). (**C,D**) I, input; U, unbound; 1, wash #1; 4, wash #4; P, IP sample. N = 3; representative blots shown.

### Activated BTK-DDX41-dependent STING signaling limits CMV protein expression and replication

Since our data indicate BTK, DDX41, and STING are activated upon lytic CMV infection, we next assessed the impact of this branch of STING signaling on infection. To this end, we utilized pharmacological inhibition using orelabrutinib, a highly specific and irreversible BTK inhibitor (40). Since BTK activates DDX41 (20), we posited this would serve as a reasonable approach to inhibit DDX41, as our attempts of both knockout and knockdown of this protein were unsuccessful (data not shown). This is perhaps not surprising, as DDX41 is embryonic lethal (30, 41). Using the pharmacological approach instead, we first evaluated the toxicity and efficacy of orelabrutinib in our model system and observed minimal toxicity over a concentration range of 0.078 to 20 µM. To validate the assay, we included puromycin treatment as a positive control (**Fig. S6A**). We also evaluated the impact of orelabrutinib treatment on the signaling pathway using a multipronged approach. We assessed DDX41 protein abundance following orelabrutinib treatment at various concentrations (0.1–10 µM) and durations (24–120 h at 0.5 µM). DDX41 abundance was attenuated following 0.5 µM orelabrutinib treatment, which was further decreased with increasing concentration of orelabrutinib (**Fig. S6B**) and longer treatment times (**Fig. S6C**). Additionally, orelabrutinib treatment reduced *IFNB* transcription in WT-infected cells, although this reduction was not statistically significant (**Fig. S6D**). Despite this, we did observe diminished phosphorylation of BTK (**Fig. S7A**) and DDX41 (**Fig. S7B**) in orelabrutinib-treated, WT-infected cells compared to untreated counterpart cultures in IP-western assays (**Fig. S7**). Our data also reveal orelabrutinib-treated, WT-infected cells display a reduction in total DDX41 abundance (**Fig. S7B**), which likely also contributes to attenuated phosphorylation levels of this protein. Collectively, these data suggest that orelabrutinib inhibit BTK and DDX41 activity without causing toxicity in fibroblasts.

Having an effective means by which to target BTK-DDX41, we leveraged orelabrutinib to assess the impact of inhibiting BTK-DDX41-dependent STING signaling on lytic CMV infection. To this end, to determine the impact of orelabrutinib-mediated DDX41 inhibition on viral protein translation, we evaluated the abundance of representative viral proteins from the early and late kinetic classes. Orelabrutinib-treated, WT-infected cells displayed a reduction in the expression of BTK, DDX41, and STING compared to infected cells treated with DMSO (− orelabrutinib; **Fig 7A**). Concurrent with the reduced abundance of these restriction factors, we observed increased abundance of the viral early protein, UL44, and the viral late protein, pp65 (**Fig. 7A**). In parallel, we evaluated CMV genome copy number in WT-infected fibroblasts treated with 0.5 µM orelabrutinib or DMSO (− orelabrutinib) over a 96 h time course. We find a statistically significant increase of viral genomes at 72 hpi in orelabrutinib-treated cells (**Fig. 7B**). We also evaluated multistep viral growth over a 16-day time course, which revealed orelabrutinib treatment resulted in an increase in cell-free virus production from 8 – 16 dpi, with a statistically significant increase at 12 dpi (**Fig. 7C**). Additionally, cell-associated virus also increased in the presence of orelabrutinib, although not to statistically significant levels (**Fig. 7D**). In sum, these data show that attenuation of BTK-DDX41-dependent signaling promotes CMV lytic replication, indicating a role for this branch of STING signaling during CMV infection.

**Figure 7.**
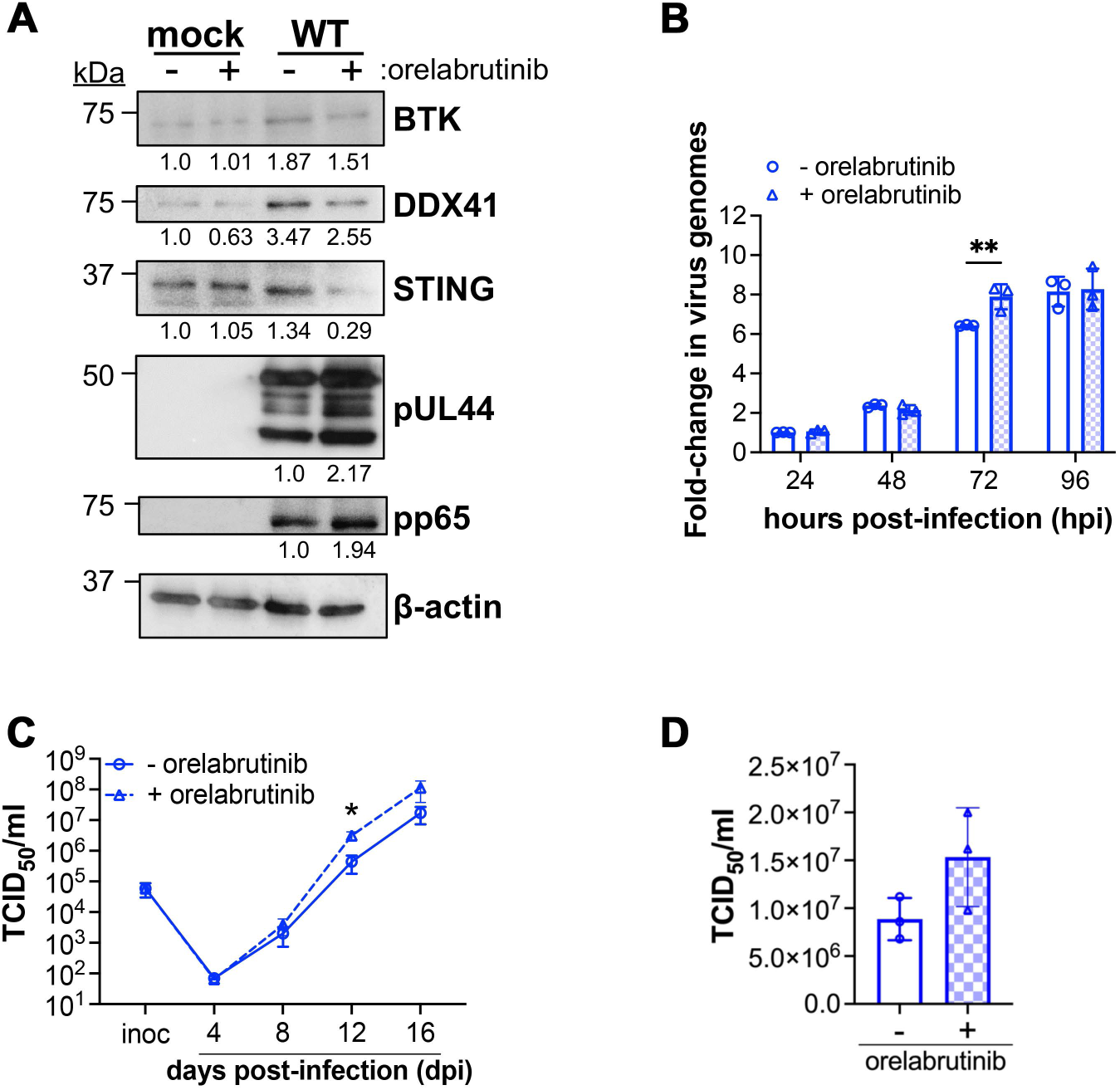
Orelabrutinib inhibition of BTK/DDX41 signaling induces lytic CMV infection. (**A, B**) NuFF-1 fibroblasts were pre-treated with 0.5 µM orelabrutinib (+ orelabrutinib) or DMSO (v/v; − orelabrutinib) for 1 h prior to mock or WT infection (MOI = 0.5 TCID_50_/cell). (**A**) At 96 hpi, cells were harvested and lysates (25 µg) probed for the indicated proteins. N = 3; representative blots shown. (**B**) Cells were harvested and whole cell DNA was isolated at the indicated time-points. Viral and cellular DNA was quantified by qPCR using primers targeting the CMV UL69 non-promoter region and cellular MDM2, respectively. The ratio of UL69 to MDM2 is shown as arbitrary units (AU) and depicted as fold-change relative to the − orelabrutinib condition at 24 h. Data represent mean values from three biological replicates, each with three technical replicates. Error bars indicate SD of three biological replicates. Statistical significance was calculated using two-way ANOVA with Tukey’s multiple comparisons test. Data are not significant unless otherwise noted; \**p* < 0.01. (**C, D**) NuFF-1 fibroblasts were treated with 0.5 µM orelabrutinib (+ orelabrutinib; open triangles/dashed line) or DMSO (v/v; − orelabrutinib; open circles/solid line) for 1 h, and then infected with WT (MOI = 0.01 TCID_50_/cell). Following adsorption, cells were washed and replenished with fresh media containing orelabrutinib (0.5 µM) or DMSO (v/v). (**C**) Cell-free virus was collected over a 16-day time course (− orelabrutinib, solid line; + orelabrutinib, dashed line), and (**D**) cell-associated virus was harvested at 16 dpi (− orelabrutinib, open bar; + orelabrutinib, checked bar). (**C,D**) Viral titers were quantified by TCID_50_ using naïve NuFF-1 fibroblasts. Samples within a biological replicate were analyzed in triplicate. N = 3; representative experiment shown. Data are mean±s.d. of the technical replicates (datapoints). Statistical significance was calculated by paired t-test. Data are not significant unless otherwise noted; \**p* < 0.05.

## Discussion

Here we show BTK-DDX41-dependent STING signaling functions in the innate immune response against lytic CMV infection. Specifically, our data reveal this branch of STING signaling is activated through the interaction of BTK and DDX41 with STING (**Figs. 1-3)**. Further, CMV infection induces a cytoplasmic redistribution of DDX41 and its associated restriction factors to the perinuclear vAC (**Fig. 4**), where it exhibits reduced activation, as evidenced by decreased tyrosine phosphorylation (**Fig. 5**) and interacts with the CMV tegument proteins pp65 and pp71 (**Fig. 6**). Additionally, we demonstrate the protective nature of this signaling pathways by pharmacological inhibition of BTK using orelabrutinib, an inhibitor that attenuates this signaling axis and results in increased viral protein abundance and CMV replication (**Fig. 7**). Collectively, our findings underscore the significance of this signaling pathway and reveal a mechanism by which CMV evades the immune response (**Fig. 8**).

**Figure 8.**
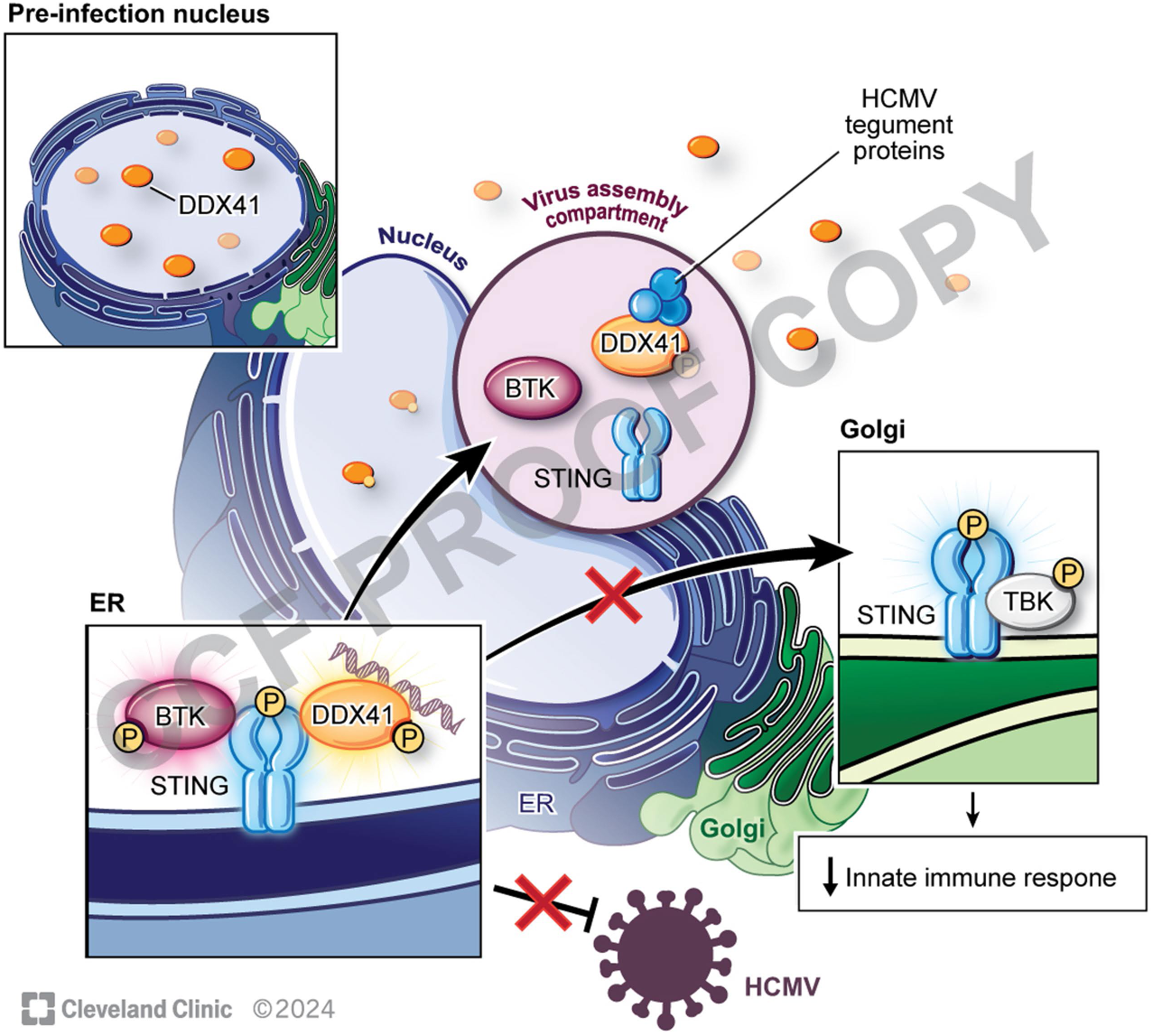
Model depicting displacement of host restriction factors during CMV lytic infection, mitigating the protective DDX41/BTK/STING signaling axis. BTK/DDX41/STING signaling is protective against lytic CMV infection. Viral infection results in displacement of the restriction factors to the virus assembly complex, where DDX41 interacts with CMV tegument proteins. Cytoplasmic redistribution of these host factors results in their decreased activity that is otherwise important for initiating antiviral signaling.

dsDNA sensors are of considerable interest in the context of CMV, as several, including cGAS, IFI16, AIM2, and ZBP1, trigger IFN production upon detection of the CMV genome (42–45). Our findings herein now add DDX41 to this growing repertoire of sensors that are involved in the innate response against CMV. Our study shows that lytic CMV infection induces the abundance of BTK, DDX41, STING, and TBK1, as well as accumulation of *IFNB* transcripts. We confirmed the activation of the pathway, as evidenced by the phosphorylation of BTK, DDX41 and STING, with both BTK and DDX41 colocalizing and interacting with STING. Previous research showed DDX41 interacts with STING at its second to fourth transmembrane domains in poly(dA)-stimulated myeloid dendritic cells (29). Additionally, in a HEK293T-based overexpression system, the BTK SH3/SH2-interaction domain binds the transmembrane region of STING (20). This mediatory role of STING is well-documented; for example, STING interacts with TBK1 and IRF3, and then facilitates the phosphorylation and activation of IRF3 by TBK1 (36). We hypothesize a similar mechanism based on the results from our study - after BTK and DDX41 bind STING, BTK could activate DDX41 through phosphorylation. Although it remains unclear whether STING directly induces this phosphorylation or merely facilitates the proximity of BTK and DDX41, current data suggest STING likely plays a more active role in the process.

Multiple studies have highlighted CMV’s diverse mechanisms to counteract DNA sensor-dependent immunity at various levels. At the DNA sensing level, CMV UL31 interacts with cGAS, causing the dissociation of DNA from cGAS and negatively regulating the STING signaling pathway (11). Similarly, UL42 binds cGAS, in turn inhibiting its DNA binding, oligomerization, and enzymatic activity (12). The tegument protein, pp65, also binds cGAS, preventing its interaction with STING and thus inactivating STING signaling (10). pp65 also interacts with IFI16’s PYRIN domain, blocking its oligomerization upon DNA sensing and subsequent immune signaling (14). Here, we observed the formation of the nascent vAC using pp65 beginning at 24 hpi - the same time point at which DDX41 was detected in the cytoplasmic fraction. This observation, as well as the documented propensity of viral proteins, particularly pp65, to bind dsDNA sensors, drew our attention to tegument proteins and their potential interaction with DDX41. Indeed, we found DDX41 associates with both pp65 and, to a lesser extent, pp71 during lytic CMV infection. The decreased detection of pp71 does not necessarily imply a weaker or less probable interaction, but rather could result from diminished availability of pp71 for interaction with DDX41 compared to pp65.

What remains outstanding is whether the interaction between DDX41 and pp65 and/or pp71 is required for relocalizing DDX41 to the vAC or whether these interactions serve to keep DDX41 in the vAC once DDX41 is already there. While possible, it is unlikely the tegument proteins aid in DDX41 redistribution. Others have reported DDX41 relocalization to perinuclear regions in response to DNA stimuli (19, 46), which Singh and colleagues demonstrated is dependent upon acetylation of DDX41’s most amino-terminal nuclear localization signal (NLS; (19)). It is thus likely that DDX41 cytoplasmic localization during CMV infection is stimulated in a similar fashion, dependent upon DNA sensing, as opposed to specific, CMV-encoded factors. Why would CMV-encoded tegument proteins interact with vAC-localized DDX41? It is possible the tegument proteins interact with DDX41 to retain this host protein in a less active state late in infection. We showed cytoplasmic DDX41 activity is attenuated, despite its colocalization with STING and BTK in the perinuclear region. It is attractive to hypothesize that the interaction with the tegument proteins precludes DDX41 activation, similar to what is reported for cGAS and IFI16 (10, 14). Alternatively, perhaps the CMV tegument protein(s)-DDX41 interaction results in DDX41’s incorporation into the mature viral particle, similar to that reported for IFI16 (18). This could ensure DDX41 is delivered to newly infected cells, allowing for its immediate action for very early innate responses following infection. This latter hypothesis could support the basis for the more late-phase phenotypes we observed herein. Further, virion-delivered DDX41 may indicate additional functions beyond innate immune responses, similar to IFI16, which also interacts with pp65 to activate viral gene transcription at the major immediate early promoter (14). DDX41 may have additional functions during CMV infection, as this host protein additionally coordinates RNA splicing and transcriptional elongation, thereby preventing DNA replication stress (47). Thus, one could posit DDX41’s functions during CMV infection are more extensive, raising the possibility its potential inclusion in the virion serves a biologically important purpose. These are compelling questions warranting further exploration, and experiments to address these possibilities are on-going.

Our findings herein highlight the protective role of BTK-DDX41-dependent STING signaling in the immune response to CMV lytic infection and further reveals a mechanism of immune evasion through disrupted activation and localization of DDX41. Our work advances our understanding of innate responses against CMV, providing insights that could inform strategies to counteract CMV’s immune evasive strategies.

## Acknowledgements

The authors thank Ian Groves for helpful discussions and Ganes Sen for critical reading of the manuscript. The authors also thank Amanda Mendelsohn (Cleveland Clinic Enterprise Creative Services) for assistance with the graphical image.

