## Supplemental Files for "The BTK-DDX41 axis of the STING pathway is activated during cytomegalovirus lytic infection"

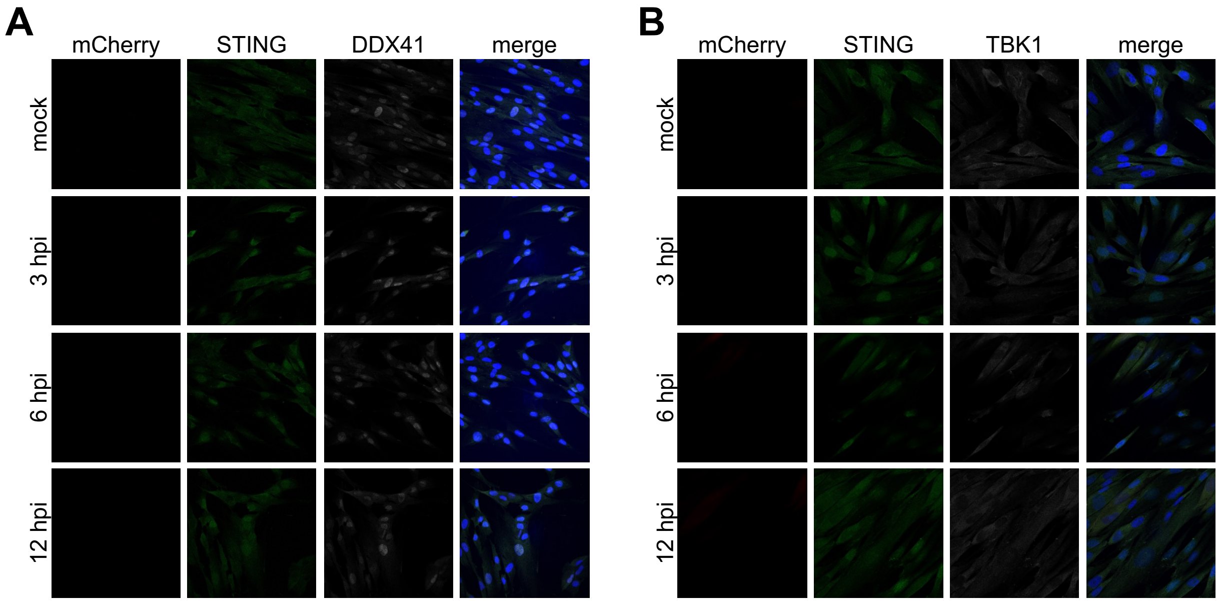


**Figure S1. Early time points for the time course in Fig. 2C,D.** NuFF-1 fibroblasts were mock- or WT-infected (MOI = 0.5 $TCID_{50}$/cell). At the indicated times post-infection, cells were processed as in Figure 2, and stained for (**A,B**) STING (each shown in green) and (**A**) DDX41 or (**B**) TBK1 (each shown in white). mCherry (red) is a marker of lytic replication and DAPI (blue) was used to visualize nuclei. Images were acquired using 63x objective. N = 3; representative images shown. As these data represent the earlier time points for the time course shown in Fig 2C,D, the mock-infected images are the same as those in the main figure.

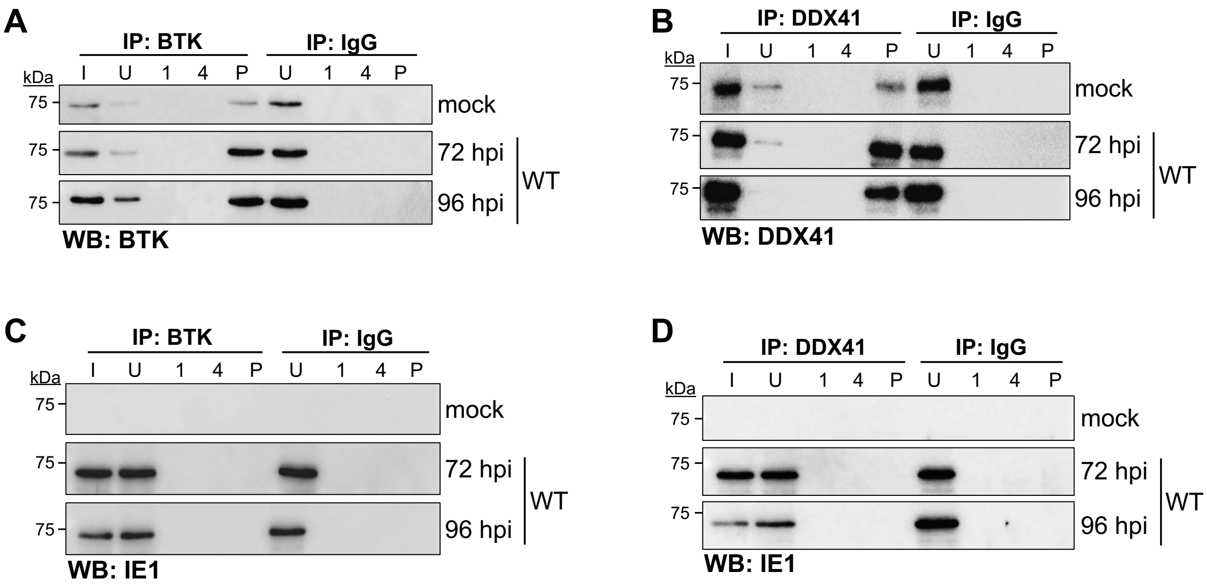


**Figure S2. Control data for findings in Fig. 3.** (**A-D**) NuFF-1 fibroblasts were mock- or WT-infected (MOI = 0.5 TCID50/cell) for the indicated times. Cells were then harvested, and immunoprecipitated (IP) with antibodies specific for (**A,C**) BTK, (**B,D**) DDX41, or (**A-D**) IgG as control. Immunoprecipitates were then probed for (**A,B**) the same antibody used for the pulldown by western blot (WB) as controls, or (**C,D**) IE1 as a marker of infection. (**A-D**) I, input; U, unbound; 1, wash #1; 4, wash #4; P, IP sample. N = 3; representative blots shown.

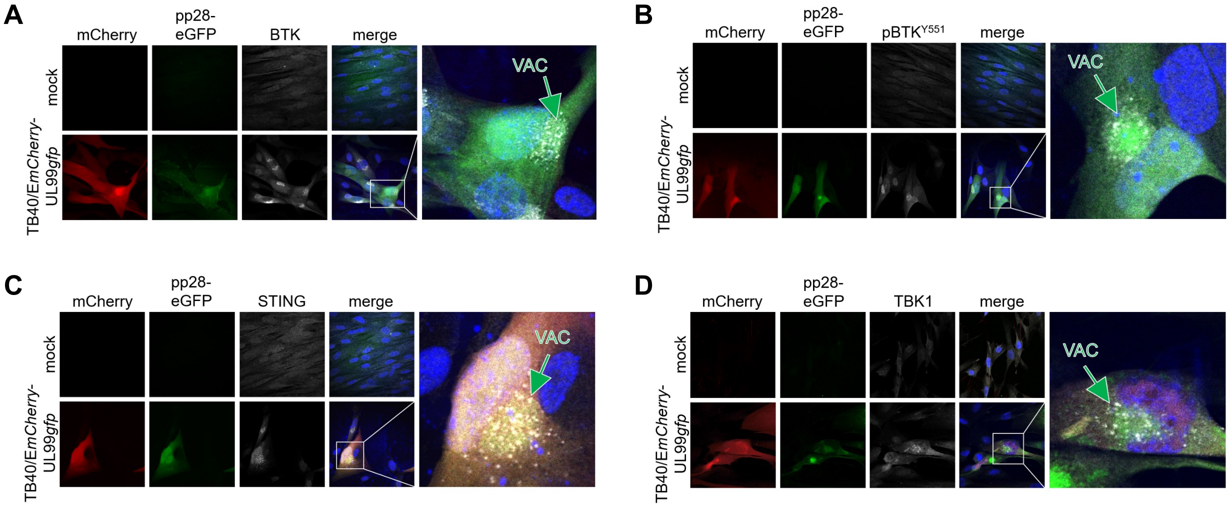


**Figure S3. DDX41-associated restriction factors localize to the viral assembly compartment (vAC).** NuFF-1 fibroblasts were mock- or TB40/E*mCherry*-UL99*gfp* (UL99*gfp*; MOI = 0.5 $TCID_{50}$/cell) for 96 h. Cells were then fixed, permeabilized, and probed for: (**A**) BTK, (**B**) $pBTK^{Y551}$, (**C**) STING or (**D**) TBK1 (each shown in white). (**A-D**) mCherry (red) is a marker of lytic infection, pp28-eGFP is shown in green, and nuclei were visualized with DAPI (blue). The viral assembly compartment (VAC) is denoted by the arrow. Images were acquired using a 63x objective with an inverted Leica SP8 confocal microscope. N = 3; representative images shown.

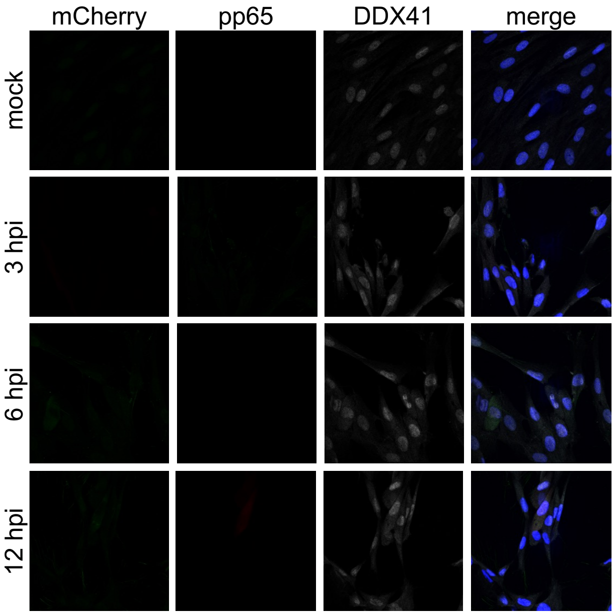


**Figure S4. Early time points for the time course in Fig. 4C.** NuFF-1 fibroblasts were mock- or WT-infected (MOI = 0.5 $TCID_{50}$/cell). At the indicated times post-infection, cells were processed as in Fig. 4, and stained for DDX41 (white) and pp65 (green). mCherry (red) is a marker of lytic replication and DAPI (blue) was used to visualize nuclei. Images were acquired using 63x objective. N = 3; representative images shown. As these data represent the earlier time points for the time course shown in Fig. 4C, the mock-infected images are the same as those in the main figure.

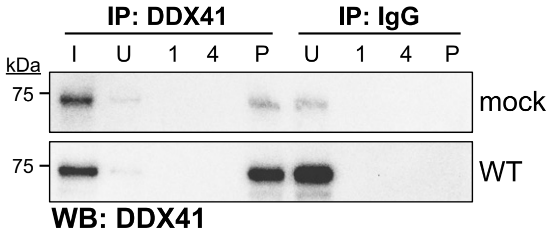


**Figure S5. Control data for Figure 6.** NuFF-1 fibroblasts were mock- or WT-infected (MOI = 0.5 TCID50/cell) for 96 h. Cells were then harvested and immunoprecipitated (IP) with antibodies specific DDX41 or IgG as a control. Immunoprecipitates were then probed for DDX41 by western blot (WB). I, input; U, unbound; 1, wash #1; 4, wash #4; P, IP sample. N = 3; representative blots shown.

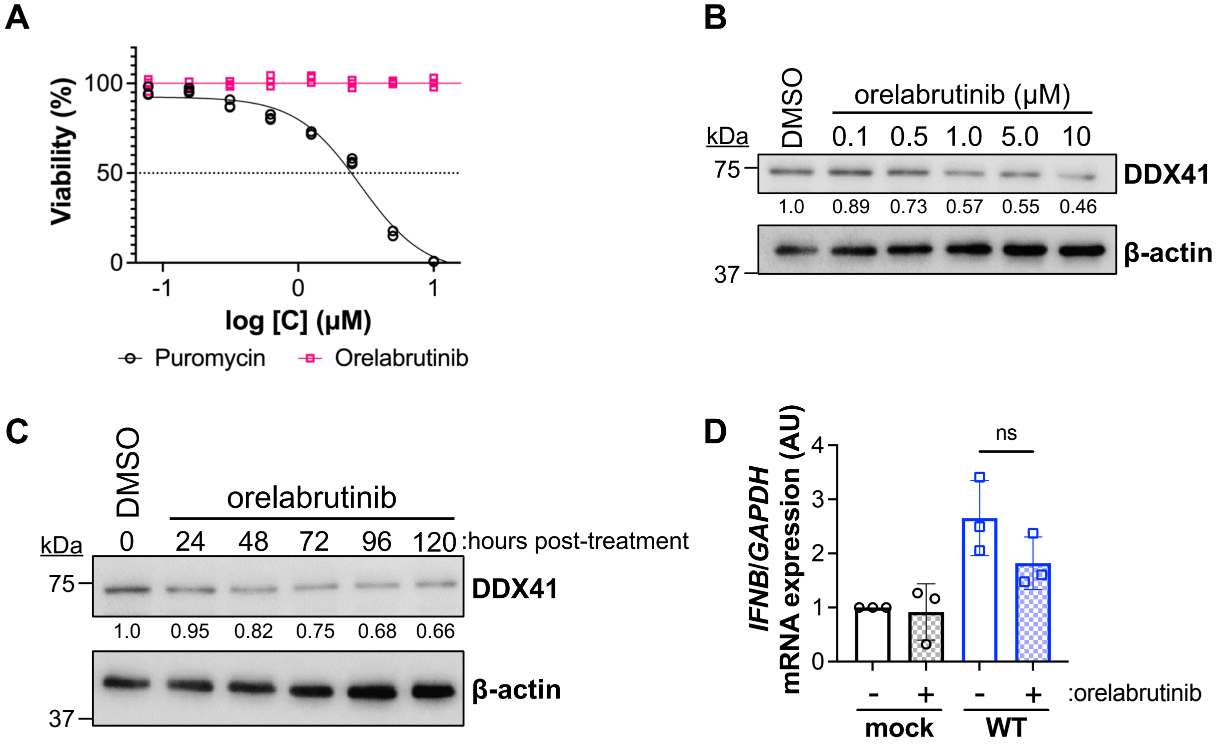


**Figure S6. The BTK inhibitor, orelabrutinib, attenuates DDX41 abundance and *IFNB* expression, with no impact on cell viability.** (**A**) NuFF-1 fibroblasts were incubated with orelabrutinib (pink; 0.08 μM – 20 μM) for 96 h. Counterpart cultures were treated with puromycin (black; 0.08 μM – 20 μM) as a positive control. Cell viability was assessed by CellTiter-Glo assay. Each concentration within a biological replicate was assessed in triplicate (indicated by datapoints). N = 3; representative assay shown. (**B**) NuFF-1 fibroblasts were treated with either orelabrutinib (0.1 – 10 μM) or DMSO (v/v). After 96 h, cells were harvested, and cell lysates (25 μg) were probed for DDX41 and cellular actin. N = 3; representative blots shown. (**C**) NuFF-1 fibroblasts were treated with 0.5 μM orelabrutinib or DMSO (v/v). Cells were harvested at the indicated times post-treatment and cell lysates (25 μg) were probed for DDX41 and cellular actin. (**B,C**) Abundance of DDX41 relative to β-actin was quantified by densitometry, relative to the DMSO control. N = 3; representative blots shown. (**D**) NuFF-1 fibroblasts were pre-treated with 0.5 μM orelabrutinib or DMSO (v/v) 1 h prior to mock (black) or WT (blue) infection (MOI=0.5 $TCID_{50}$/cell). Cells were harvested 96 hpi, total RNA was isolated, and *IFNB* mRNA expression was quantified by RT-qPCR. Data is plotted as ΔΔCt for *IFNB* relative to *GAPDH* in arbitrary units (AU). Each sample within a biological replicate was analyzed in triplicate (datapoints). Error bars indicate the SD of the mean of three biological replicates. ns, not significant. N = 3; representative assay shown.

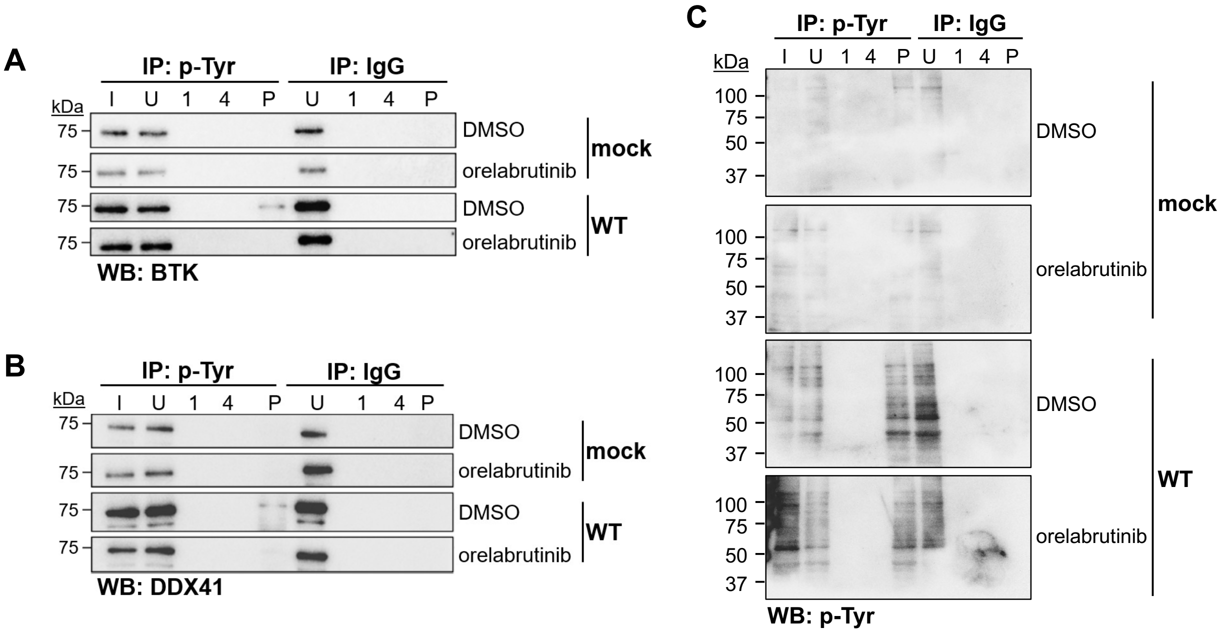


**Figure S7. Orelabrutinib inhibits CMV-induced DDX41 and BTK phosphorylation.** NuFF-1 fibroblasts were pre-treated with 0.5 μM orelabrutinib or DMSO (v/v) 1 h prior mock or WT infection (MOI = 0.5 $TCID_{50}$/cell) for 96 h. Cells lysates were immunoprecipitated (IP) with antibodies specific for phosphorylated-tyrosine (p-Tyr) or IgG. Immunoprecipitates were then probed for (**A**) BTK, (**B**) DDX41, or (**C**) p-Tyr by western blot (WB). I, input; U, unbound; 1, wash #1; 4, wash #4; P, IP sample. N = 3; representative blots shown.
